# Size Control of hnRNPK-based Nucleolar Condensates by RNA-Regulated Fusion Dynamics

**DOI:** 10.64898/2026.09.14.751346

**Authors:** Andrés R. Tejedor, Juan Luengo, Pablo Llombart, Eduardo Pedraza, Álvaro Otero-Sobrino, Maria Velasco-Estevez, Miguel Gallardo, Alberto Ocana, Rosana Collepardo, Jorge R. Espinosa

## Abstract

Nucleoli are liquid-like condensates whose size is actively conserved—they remain small, numerous, and resistant to coalescence—yet the molecular mechanisms that constrain their fusion remain poorly understood. Heterogeneous nuclear ribonucleoprotein K (hnRNPK), an RNA-binding protein implicated in nucleolar organization and cancer, interacts directly with the scaffold protein Nucleolin. Combining residue-resolution coarse-grained simulations with biochemical experiments, we find that hnRNPK and Nucleolin condense through distinct interaction networks—a localized cation–*π*/electrostatic hotspot in hnRNPK versus broadly distributed electrostatic contacts in Nucleolin, reorganized upon co-assembly. To probe how RNAs reshape these condensates, we develop and validate, against re-entrant phase-separation experiments and AlphaLISA binding data, a nucleotide-resolution coarse-grained model for single-stranded RNA. Using this framework, we show that RNA is asymmetrically and preferentially recruited by hnRNPK over Nucleolin, an asymmetry that grows stronger when the two proteins compete for the same RNAs. This selective recruitment sustains a dynamic fission-fusion equilibrium: hnRNPK-containing condensates repeatedly fuse and split rather than coalescing into a single condensate, whereas Nucleolin-containing and ternary condensates fuse into one dominant cluster. These results reveal a molecular mechanism, grounded in sequence-encoded, RNA-controlled fusion dynamics, by which nucleolar condensates conserve a controlled, non-coalescing size despite their liquid-like character.

## I. INTRODUCTION

The nucleolus is the largest and most extensively studied biomolecular condensate in the eukaryotic nucleus, serving as the principal site of ribosomal RNA (rRNA) transcription, processing, and pre-ribosome assembly^1–3^. Rather than being enclosed by a membrane, it assembles through liquid–liquid phase separation (LLPS) into a multiphase, multilayered condensate whose immiscible sub-compartments spatially organize the sequential steps of ribosome biogenesis^4–6^. Yet unlike a simple liquid droplet, nucleoli typically remain small and numerous within a single nucleus rather than coalescing into one large body, a characteristic size actively maintained through concentration-dependent phase behaviour and controlled microdroplet fusion^7,8^. This size control is not absolute: nucleolar number, size, and shape vary systematically with cellular state, becoming smaller under conditions that extend organismal lifespan^9^, while nucleolar hypertrophy, an increased number of nucleoli, and fragmentation into disorganized structures are recurrent hallmarks of cancer, ribosomopathies, and ageing^4,10^.

Among the molecular regulators of nucleolar organization and size, heterogeneous nuclear ribonucleoprotein K (hnRNPK) stands out as a multifunctional RNA-binding protein (RBP) with central roles in gene regulation, mRNA processing, and ribosome biogenesis^11–13^. hnRNPK localizes to the nucleolus, where it interacts directly with Nucleolin^11,14^, a major scaffold protein that regulates ribosomal RNA synthesis, pre-rRNA processing, and the assembly, internal organization, and material properties of nucleolar condensates^15,16^. Its physiological importance is illustrated by the fact that hnRNPK overexpression induces p53-dependent nucleolar stress, leading to cell-cycle arrest, senescence, and bone marrow failure phenotypes reminiscent of ribosomopathies^11^. This close functional and physical association with Nucleolin raises several fundamental questions. Do the two proteins rely on common or distinct molecular mechanisms to drive condensate formation? How are these mechanisms remodelled upon heterotypic RNA-driven co-assembly? And, ultimately, how do they contribute to keeping nucleolar condensates small and resistant to coalescence? Understanding how hnRNPK and Nucleolin establish and reorganize their intermolecular interaction networks is therefore essential for elucidating how nucleolar condensates assemble, maintain their architecture, and regulate their biological function.

Moreover, far beyond the dynamic organization of the nucleolus, LLPS is recognized as a universal mechanism by which cells compartmentalize proteins and nucleic acids into dynamic membraneless condensates throughout the nucleus and the cytoplasm^17–22^. The assembly of these condensates arises from large networks of weak, multivalent interactions encoded within the intrinsically disordered regions (IDRs) of RBPs together with their interactions with RNAs^23–25^. Consequently, subtle sequence-dependent modifications can profoundly remodel not only condensate density, viscosity, molecular composition, and stability, but also their propensity to fuse or remain as discrete, size-limited assemblies, by rewiring the underlying inter-molecular network connectivity^26–30^. Through these emergent physicochemical properties, condensates regulate fundamental nuclear processes including transcription, RNA metabolism, DNA repair, and genome organization, while their dysregulation has emerged as a common molecular hallmark of numerous cancers and neurodegenerative diseases^31–41^. However, despite their central biological importance, the molecular interaction networks that govern condensate self-assembly, co-condensation, and fusion dynamics remain largely inaccessible to experimental characterization because of the multicomponent, dynamic, heterogeneous, and densely packed nature of the condensed phases^22,42–45^.

Molecular dynamics (MD) simulations, and in particular coarse-grained models, have emerged as a powerful approach to overcome these experimental limitations, enabling the characterization of the sequence-encoded molecular interactions that govern condensate formation across size- and timescales inaccessible to atomistic simulations^46–54^. Over the past few years, several residue-resolution coarse-grained force fields have been developed to describe biomolecular condensation, including HPS-based models^46–48^, the CALVADOS family^49–51^, and the Mpipi/Mpipi-Recharged force fields^53,55,56^. Among these, the Mpipi-Recharged introduces residue pair-specific electrostatic interactions through a Yukawa potential^57,58^, providing a chemically resolved description of electrostatics that substantially improves predictive accuracy across a broad range of IDPs and multi-domain protein condensates^59–62^. This pair-specific treatment of electrostatic interactions makes the model particularly well suited to investigate condensates containing highly charged sequences such as RBPs^63,64^, where electrostatic interactions play a dominant role in determining condensate stability, composition, and material properties^64–66^. Yet a nucleotide-resolution coarse-grained model for RNA compatible with the Mpipi-Recharged, able to capture protein–RNA interactions with the same specificity, has so far been lacking, limiting quantitative investigation of how different RNAs reshape condensate assembly, organization, and dynamics.

To address these questions, we perform residue-resolution coarse-grained molecular dynamics simulations to dissect how hnRNPK and Nucleolin assemble into nucleolar condensates, individually and together. Using the Mpipi-Recharged^55^, we first map the sequence-resolved interaction networks underlying condensation of each protein and show that near-identical phase diagrams can mask markedly different underlying molecular grammars, which are further reorganized upon coassembly. We then extend this framework to protein– RNA interactions by developing and experimentally validating a nucleotide-resolution coarse-grained model for single-stranded RNA, and use it to determine how the two proteins partition a shared RNA solution. This analysis reveals that RNA engagement is markedly asymmetric between the two proteins, and that this asymmetry has direct consequences for condensate size: hnRNPK-containing assemblies persistently fragment into multiple, smaller nuclei rather than coalescing into one large condensate, whereas Nucleolin-containing and mixed condensates readily converge into a single, larger structure. Together, these results establish a molecular framework linking sequence-encoded protein and RNA interactions to the assembly, internal organization, and conserved size of nucleolar condensates.

## II. RESULTS

### A. Distinct sequence-encoded interaction networks in hnRNPK and Nucleolin drive similar condensate behaviour

Nucleolin and hnRNPK, two of the principal RBP scaffolds that co-reside within the nucleolus^11,15,16^, offer a computationally tractable system for dissecting how protein sequence composition encodes the intermolecular interactions that drive nucleolar condensate formation. Both are highly charged RBPs with different IDRs, whose sequence compositions are expected to promote biomolecular condensation through multivalent intermolecular contacts^5,67^. Despite sharing a high abundance of positively and negatively charged residues interspersed with aromatic amino acids (Fig. 1A, top), the two proteins exhibit markedly different sequence organizations. Whereas the charged residues of hnRNPK are relatively evenly distributed along the sequence, Nucleolin displays a substantially more heterogeneous charge pattern, with extended negatively charged domains alternating with clusters of positively charged residues. In particular, the N-terminal region of Nucleolin (residues ∼50–140) contains a dense positively charged segment, whereas residues ∼150–270 comprise several extended negatively charged patches that are largely absent in hnRNPK (Fig. 1A, top). Such differences in charge patterning are expected to profoundly influence the intermolecular interaction network driving condensation, potentially giving rise to distinct phase behaviours and spatial organizations characteristic of highly charged condensates64,65,68,69.

**FIG. 1.**
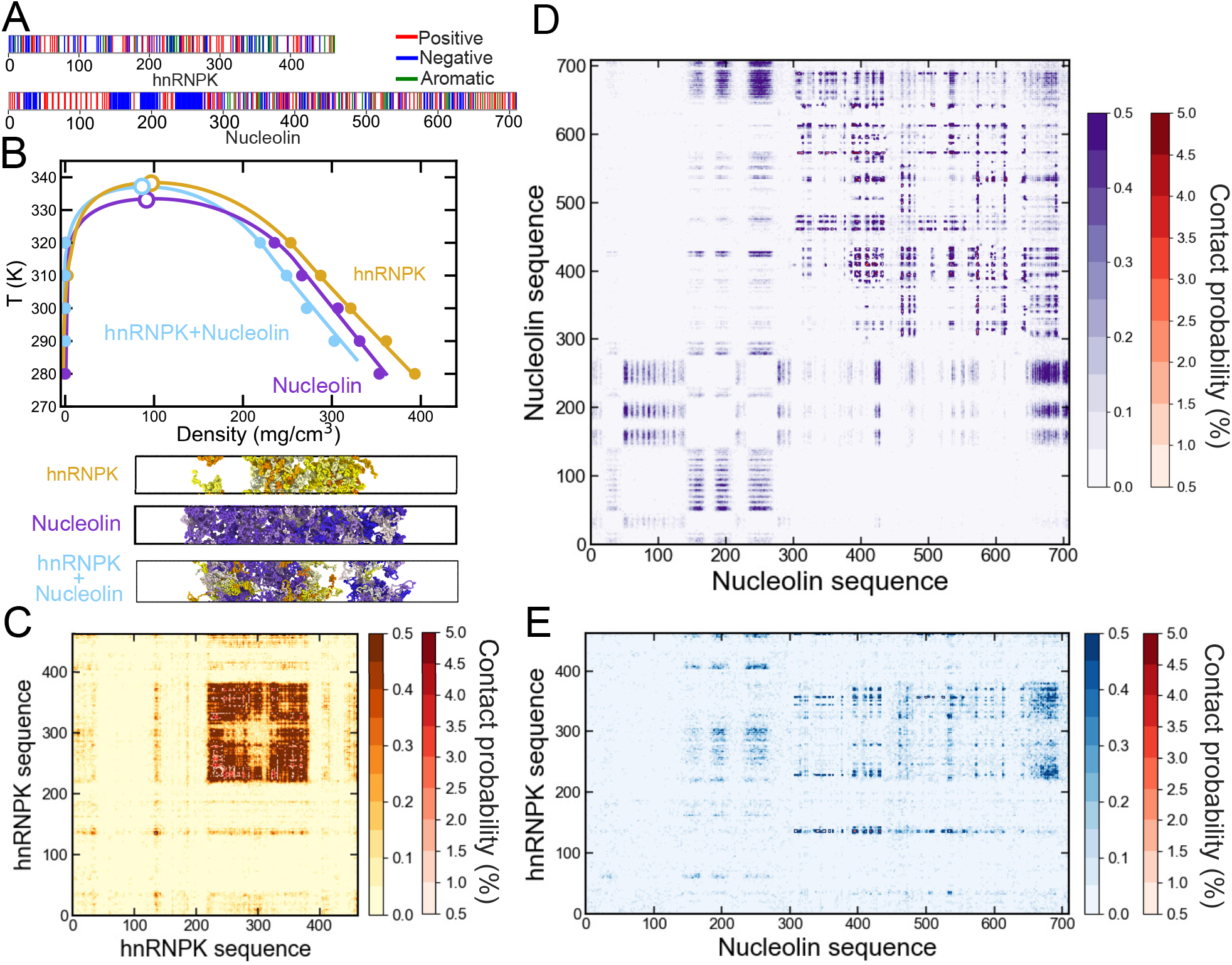
(A) Sequence composition of hnRNPK and Nucleolin highlighting positively charged, negatively charged, and aromatic residues as indicated. (B) Phase diagrams in the temperature–density plane for hnRNPK (golden), Nucleolin (purple), and the equimolar hnRNPK–Nucleolin mixture (light blue), with the critical point of each system represented by an empty symbol and the coexistence lines depicted as a guide to the eye. Representative snapshots from Direct Coexistence (DC) simulations at 310 K are displayed below for each of the three systems. (C–E) Intermolecular contact map of hnRNPK (C), Nucleolin (D), and hnRNPK-Nucleolin mixture obtained from DC simulations of the condensates at 310 K and 150 mM NaCl concentration. Colorbars indicate the contact probability in percentage, with a two-tone scale distinguishing low- and high-probability contacts.

To quantify the condensation propensity of hnRNPK and Nucleolin, we first performed Direct Coexistence (DC) simulations^70–72^ using the implicit-solvent residue-resolution Mpipi-Recharged model (see Section SI of the SM for technical details on the model potential and parameters)^55^. At subcritical temperatures, the simulations spontaneously demix into coexisting condensed and dilute phases, enabling direct determination of the coexistence densities (representative snapshots at 310 K are shown in Fig. 1B, bottom). The resulting phase diagrams (Fig. 1B, centre) reveal that hnRNPK and Nucleolin exhibit remarkably similar coexistence boundaries despite their markedly different sequence architectures and chain lengths. Strikingly, although Nucleolin is substantially longer than hnRNPK, its higher overall charge density compensates for the higher entropic cost of condensation expected for longer polymer chains, resulting in critical temperatures that differ by only ∼10 K. These critical temperatures are comparable to those previously reported for archetypal condensate-forming proteins, including FUS^61,73^, hnRNPA1^60,75^, or TDP-43^63,77^, indicating that both proteins possess a robust intrinsic propensity to undergo condensation driven by homotypic interactions.

To understand how hnRNPK and Nucleolin achieve nearly identical condensation propensities despite their markedly different sequence architectures, we next analyse the intermolecular interaction networks that stabilize their condensed phases using residue-resolved contact maps^79,80^ (see Section SIII of the SM for methodological details on these calculations). These maps quantify the probability of intermolecular contacts between every pair of residues across different protein replicas sampled in the DC simulations (Fig. 1C, D), thereby revealing the sequence domains that contribute most strongly to condensate cohesion. For hnRNPK, intermolecular contacts are highly localized within a region spanning residues ∼220–360, corresponding to the sequence flanking the KH domains that is strongly enriched in arginine and aromatic residues (Fig. 1A), consistent with deletion-construct mapping studies that identify this same region as required for hnRNPK–RNA binding^81^. As expected from its sequence composition, this hotspot is dominated by cation–*π* interactions between arginine side chains and aromatic residues, complemented by favourable electrostatic interactions, as confirmed by the decomposition of the most frequent residue-residue contacts (Fig. S1). In contrast, Nucleolin exhibits no comparable interaction hotspot. Instead, intermolecular contacts are broadly distributed throughout the sequence, indicating that condensate stability arises from a highly cooperative network of long-range electrostatic interactions established between its alternating positively and negatively charged patches (Fig. 1A). These results reveal that the similar phase behaviour of hnRNPK and Nucleolin emerges from genuinely distinct microscopic interaction networks: hnRNPK condensation is driven by a localized hotspot dominated by cation–*π* contacts, whereas Nucleolin relies on distributed electrostatic interactions spanning much of its sequence.

### B. Co-assembly rewires the Nucleolin’s interaction network and generates microphase-separated condensates

While hnRNPK and Nucleolin are known to co-localize in the nucleus, where hnRNPK directly interacts with nucleolar components including Nucleolin^13,14^, it remains unclear whether these proteins co-assemble into a common condensate and how heterotypic interactions modify the condensation mechanisms identified for the individual proteins. Motivated by this functional association, we perform Direct Coexistence simulations of an equimolar hnRNPK–Nucleolin mixture to determine its phase behaviour (Fig. 1B). Remarkably, the binary condensate exhibits a phase-separation coexistence line and critical temperature nearly identical to those of the corresponding single-component systems, differing by only a few kelvin from the pure hnRNPK and Nucleolin phase diagrams (Fig. 1B). This result indicates that heterotypic interactions neither strongly enhance nor suppress the overall thermodynamic driving force for phase separation; instead, the condensate remains primarily stabilized by the intrinsic condensation propensities of each protein. Consistent with this interpretation, representative snapshots at 310 K (Fig. 1B, bottom) reveal a micro-heterogeneous condensed phase in which hnRNPK- and Nucleolin-rich regions remain partially segregated, with only limited intermixing between the two protein species, a hallmark of condensates formed by highly charged macromolecules^64,82–84^.

To determine how co-condensation modifies the molecular interaction network, we next analyse the residue-resolved intermolecular contact maps of the binary mixture (Fig. 1E). Remarkably, the homotypic interaction networks of both proteins remain largely unchanged upon mixing. The hnRNPK–hnRNPK contact map (Fig. S2 in the SM) is virtually identical to that of the pure condensate, preserving both the localization and intensity of the characteristic interaction hotspot spanning residues ∼220–360 (Fig. 1C). Likewise, the homotypic Nucleolin contact network remains broadly distributed throughout the sequence, closely resembling that observed in the single-component condensate (Fig. S3 in the SM). These results indicate that the intrinsic molecular grammars governing condensation of each protein are remarkably robust to the presence of the second component. On the other hand, the heterotypic hnRNPK–Nucleolin contact map (Fig. 1E) displays a diffuse pattern of interactions distributed over both protein sequences: rather than forming a specific interaction hotspot or dominant binding interface, the two proteins establish numerous weak contacts across multiple regions of their sequences. This diffuse interaction pattern is fully consistent with the partially segregated architecture observed in the condensed phase, where heterotypic interactions are sufficient to maintain a shared condensate but not sufficiently strong to overcome the intrinsic homotypic interaction preferences of either protein. Consequently, co-condensation gives rise to micro-segregated condensates in which each protein largely preserves its own sequence interaction network while remaining dynamically coupled through weak and highly distributed heterotypic interactions.

Inspection of the most-frequent heterotypic contacts between hnRNPK and Nucleolin (Fig. S1 in the SM) reveals that the intermolecular interface is dominated by electrostatic favourable interactions (blue), with a substantial contribution from cation–*π* contacts (pink). This interaction pattern follows directly from the sequence architectures of the two proteins (Fig. 1A): the arginine- and aromatic-rich region of hnRNPK (residues ∼220–360), which constitutes its principal condensation hotspot, is ideally positioned to interact with the highly charged segments distributed throughout the Nucleolin sequence through complementary electrostatic and cation–*π* interactions. Moreover, particularly frequent residue contact pairs also include GD, GE, and EF contacts, indicating that the glycine-rich regions of hnRNPK establish extensive contacts with the acidic glutamate- and aspartate-rich stretches of Nucleolin, while phenylalanine residues from the hnRNPK hotspot also contribute to interactions with negatively charged glutamate residues. Together, these contacts illustrate how the complementary sequence composition of the two proteins gives rise to a broad, distributed heterotypic interface. Notably, whereas cation–*π* interactions are the principal driving force underlying hnRNPK self-association, they contribute only partially to Nucleolin condensation. Consequently, co-condensation recruits interaction modes that are less dominant from the major condensation mechanism of each of the individual components, giving rise to a distinct heterotypic interaction grammar. These observations demonstrate that condensate composition does not simply tune the relative contributions of pre-existing interactions, but can generate emergent molecular networks with physicochemical properties that are inaccessible to the isolated proteins. More generally, these results suggest that the molecular grammar of condensates is itself composition-dependent, providing cells with a versatile mechanism to expand the functional and structural diversity of condensates through combinatorial assembly of many different protein components.

### A transferable coarse-grained RNA model captures re-entrant phase behaviour and hnRNPK–RNA co-condensation

RNA is an integral component of nucleolar function as well as many other nuclear biomolecular condensates, where it regulates phase separation through a combination of electrostatic interactions and sequence-specific recognition by RNA-binding proteins^47,85,86^. Consistent with this central role, hnRNPK binds numerous coding and non-coding RNAs to regulate RNA metabolism, translation, and gene expression, with several of these interactions playing direct roles in tumorigenesis^11^. Notably, recent studies have identified hnRNPK as a primary cellular target of small molecules that suppress MYC expression by promoting the cytoplasmic relocalization of hnRNPK and the sequestration of MYC mRNA into stress granules^87–90^. This relocalization-based mechanism is mechanistically distinct from the direct binding of hnRNPK to MYC transcripts that promotes their translation^11^, and the two should not be conflated: our discussion of hnRNPK as a druggable node throughout this work refers specifically to the former, condensate-disrupting strategy. These findings establish hnRNPK–RNA interactions as not only fundamental regulators of condensate biology, but also as promising therapeutic targets. Despite their biological and pharmacological importance, the molecular mechanisms governing hnRNPK–RNA co-condensation remain poorly understood, largely because the sequence-dependent interactions between disordered RNA molecules and intrinsically disordered proteins are difficult to resolve experimentally^42^. To address this challenge, we next develop a nucleotide-resolution coarse-grained parameterisation for single-stranded RNAs fully compatible with the Mpipi-Recharged force field^55^, enabling quantitative investigation of sequence-dependent protein–RNA interactions and their role in condensate formation. The corresponding protein–RNA cross-interaction parameters were derived using a bottom-up strategy based on potentials of mean force (PMF) calculated from all-atom MD simulations performed by us^91^. Consistent with the original force field, short-range interactions are described using the Wang–Frenkel potential, whereas long-range electrostatic interactions are represented through screened Yukawa potentials. This unified parameterisation enables proteins and disordered RNAs to be simulated within a common molecular resolution framework while preserving the physicochemical consistency of the original Mpipi-Recharged model. The complete set of interaction parameters and potentials for the RNA model is provided in Sections SI and SIV of the SM.

To assess the predictive capability of the proposed protein–RNA interactions, we benchmark the RNA model against the re-entrant phase-separation experiments of Alshareedah *et al*.^26^. In these experiments, turbidity was measured as a function of the RNA/peptide mass ratio for mixtures of polyuridylic acid (polyU) or polyadenylic acid (polyA) with arginine-rich (R) and lysine-rich (K) peptides. Upon RNA addition, turbidity initially increases as electrostatic and cation–*π* attractions promote condensate formation, reaches a maximum at intermediate RNA concentrations (near the electroneutral point), and subsequently decreases at excess RNA concentration due to re-entrant dissolution driven by overcharging of the condensates and the resulting long-range electrostatic RNA–RNA self-repulsion^26,92^. This characteristic bell-shaped phase behaviour provides a stringent benchmark for evaluating whether a coarse-grained model can simultaneously capture both the attractive and repulsive regimes governing protein–RNA complex coacervation.

To reproduce these experiments computationally, we perform bulk NpT simulations—as proposed in Refs.^97,98^—at *p* = 0 bar over a broad range of RNA/peptide compositions and temperatures. Condensate formation is identified from the relaxed density of the system, with condensed states defined as those with *ρ* ≳ 0.1 g cm^−397,99^. The corresponding critical solution temperature for each system is bracketed between the highest temperature sustaining a stable condensed phase and the lowest temperature at which complete dissolution occurs. In Figure 2 we compare the predicted phase-separation behaviour (Fig. 2B), representative simulation snapshots (Fig. 2A), and the experimentally measured turbidity curves from Ref.^26^ normalized by their maximum intensity (Fig. 2C). Notably, in Fig. 2B, while points near the maximum in 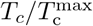 correspond to liquid-like condensates, our simulations of RNA-peptide ratios far away from charge compensation instead result in the formation of more kinetically arrested complexes. Our simulations qualitatively reproduce the characteristic re-entrant phase behaviour of all four RNA–peptide systems, with the condensates formed by arginine-rich peptides and polyA RNAs displaying higher stability. In every case, the normalized critical temperature, 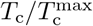, exhibits the same bell-shaped dependence on RNA/peptide mass ratios observed experimentally, demonstrating that the model correctly captures the competition between short-range attractive interactions that stabilize condensates at intermediate RNA concentrations and long-range electrostatic repulsion that drives their dissolution at RNA excess^92,93,100^.

**FIG. 2.**
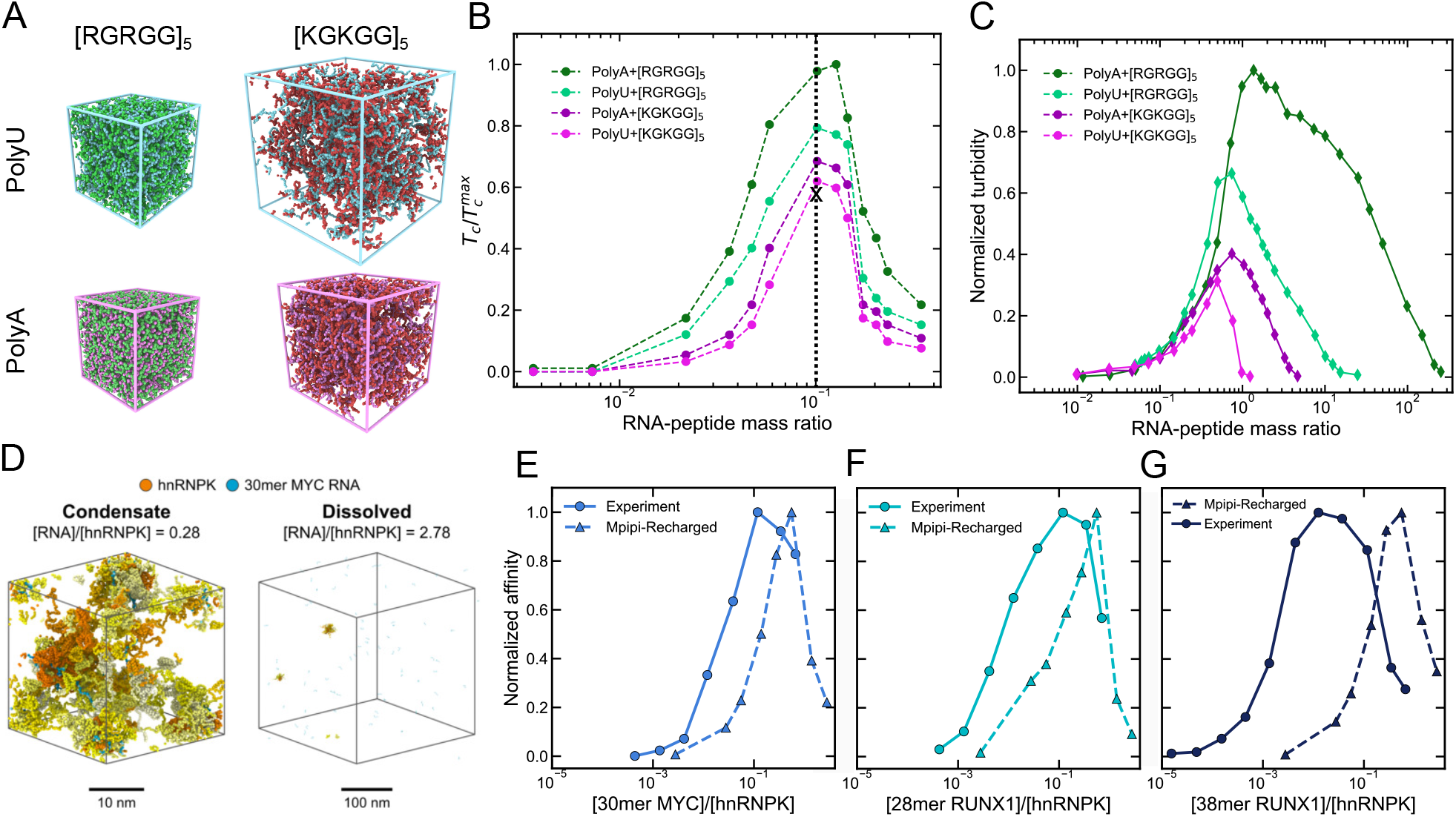
(A) Representative NpT simulation snapshots for the four RNA–peptide systems used to validate the coarse-grained RNA model. PolyU systems are shown in the top row and PolyA systems in the bottom row; the arginine-rich peptide [RGRGG]5 is shown in the left column and the lysine-rich peptide [KGKGG]5 in the right column. RNA chains are coloured according to identity (PolyU in cyan, PolyA in pink) and peptides according to identity (green for [RGRGG]5, red for [KGKGG]5). (B) Normalized 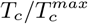 from NpT simulations as a function of the RNA/peptide mass ratio for all four systems. The dashed vertical line and cross symbol mark the reference ratio and temperature used for the snapshots in panel A. (C) Corresponding normalized turbidity curves from the experiments of Alshareedah *et al*. (2019) for the same four systems. (D) Representative NpT simulation snapshots at 300 K of hnRNPK in the presence of 30mer MYC RNA at two different RNA-to-protein molar ratios, illustrating the reentrant phase behaviour: a condensed state (left, [RNA]*/*[hnRNPK] = 0.28) and a dissolved state (right, [RNA]*/*[hnRNPK] = 2.78). hnRNPK protein replicas are shown in orange and RNA strands in cyan. Scale bars are 10 nm (left) and 100 nm (right). (E–G) Interaction profiles between hnRNPK (kept constant at 7.5 *µ*g/ml) and increasing concentrations of (**E**) 30mer MYC DNA, (**F**) 28mer RUNX1 DNA, and (**G**) 38mer RUNX1 DNA as measured by AlphaLISA. The x-axis reports the molar ratio of DNA to hnRNPK and the y-axis (left) reports the normalized affinity (mean ±SD, *n* = 3). Simulated intermolecular contact frequencies (y-axis; right) from condensate simulations at 300 K are overlaid as dashed lines.

We find that the agreement remains robust across both polyA and polyU RNAs and for peptides enriched in either arginine or lysine, indicating that the parameterization is transferable across chemically distinct protein– RNA systems. Nevertheless, our simulations underestimate the RNA concentration at which re-entrant dissolution sets in, with the predicted critical temperature beginning to decrease at lower RNA/peptide ratios than observed experimentally for turbidity (Fig. 2C). This discrepancy does not reflect a shortcoming of the underlying interaction parameters; rather, it stems from a computational constraint. Experimentally, polyU and polyA RNA strands range from approximately 1500 to 3000 nucleotides in length^26^, whereas in our simulations RNAs are 200 nucleotides long. As shown in Ref.^99^, condensate stability increases with RNA length, and the RNA/peptide ratio at which stability is maximal shifts toward much higher RNA concentrations as RNA length increases. Reproducing this effect for 1500– 3000-nucleotide strands would require a correspondingly larger simulation box to prevent RNA chains from self-interacting across the periodic boundary conditions—a system size that cannot be sampled within a reasonable simulation time. Hence, the discrepancy in the concentration of re-entrant dissolution is primarily a consequence of this length-dependent finite-size constraint, rather than of the model itself. Because this constraint only becomes limiting as RNA length approaches the simulation box dimensions, comparisons for RNAs at or below the ∼200-nucleotide length are not subject to this computational limitation. Furthermore, as a second-order approximation, the present RNA model does not explicitly account for sequence-dependent base-stacking interactions, ion-mediated RNA–RNA interactions, and secondary-structure formation, all of which may further reduce electrostatic self-repulsion under highly RNA-rich conditions^101–103^. Nevertheless, despite these coarse-grained approximations, the model is able to reproduce the experimentally observed transition from RNA-induced condensate stabilization to re-entrant dissolution, providing a transferable framework for investigating sequence-dependent protein–RNA condensates.

Having validated the protein–RNA force field against re-entrant phase-separation experiments, we next apply it to investigate the interaction of hnRNPK with biologically relevant nucleic acid sequences associated with its transcriptional targets. Specifically, we study hnRNPK binding to three single-stranded RNA elements derived from the promoter regions of *RUNX1* (28-mer and 38-mer) and *MYC* (30-mer; see further details on the sequences in Section SII of the SM)—two key oncogenic regulators whose expression is controlled by hnRNPK in hematological malignancies^12,81,87,89,90,104^. We perform simulations under NpT conditions at 300 K over a wide range of RNA concentrations to characterize the sequence-dependent interaction profiles of hnRNPK. The resulting normalized hnRNPK–RNA contact frequency as a function of nucleic acid concentration is then qualitatively compared with AlphaLISA DNA binding measurements, in which the hnRNPK concentration was held constant at 7.5 *µ*g ml^−1^ while the concentration of each DNA construct was systematically titrated (Fig. 2E–G; full experimental details are provided in Section SVI of the SM).

Both the simulations and the AlphaLISA experiments reveal a clear concentration-dependent interaction between hnRNPK and the RUNX1 and MYC nucleic acid constructs (Fig. 2E–G). In the simulations, varying the nucleic acid-to-protein ratio induces pronounced changes in the protein–RNA contact frequency and condensate density, which directly modulate the intermolecular network connectivity and, consequently, the probability of hnRNPK–nucleic acid interactions (Fig. 2E– G). Experimentally, the AlphaLISA signal increases with DNA concentration, reaches a maximum at intermediate DNA/protein ratios, and decreases at DNA excess, consistent with the characteristic stoichiometric hook effect arising from saturation of hnRNPK high-affinity binding sites. Remarkably, the normalized contact frequencies extracted from our simulations closely reproduce this bell-shaped concentration dependence for both the RUNX1 and MYC constructs, demonstrating that our coarse-grained model captures the sequence-dependent recognition of biologically relevant hnRNPK targets.

We note that, although our force field was parameterized and validated using RNA systems (Fig. 2), its benchmark against protein–DNA complexes is physically grounded since the interaction landscape of hnRNPK is mostly dominated by long-range electrostatic interactions between its highly charged IDRs and the negatively charged phosphodiester backbone common to all nucleic acids, complemented by local cation–*π* interactions involving aromatic and positively charged residues effectively integrated into our model through Wang–Frenkel dispersive interactions. These physicochemical molecular interactions are largely shared between RNA and DNA, making the resulting binding mechanism largely independent of the identity of the nucleic acid polymer as long as the sequence composition and length are preserved. Importantly, this agreement is maintained—and is even more compelling—in the present biologically realistic benchmark, involving full-length hnRNPK and (28–38)-mer nucleic acid constructs. In contrast to the simplified peptide–RNA systems employed for force field validation (Fig. 2B–C), this comparison probes molecular systems whose sizes and sequence complexity are much closer to those encountered *in vitro*^26^, providing a substantially more stringent assessment of our model’s predictive capability. Consequently, the agreement between simulations and experiments for the three nucleic acid sequences demonstrates that the model captures the underlying physicochemical grammar governing hnRNPK–nucleic acid recognition rather than merely reproducing RNA-specific phase behaviour. Taken together, these results establish that the Mpipi-Recharged framework, extended here to single-stranded RNAs, accurately predicts concentration-dependent, sequence-sensitive recognition of biologically relevant nucleic acid targets. More broadly, they show that residue-resolution coarse-grained simulations provide a transferable computational framework for investigating protein–nucleic acid interactions, opening the possibility of large-scale *in silico* screening of multicomponent hnRNPK target sequences and rationally predicting how sequence variation, post-translational modifications, condensate composition, or disease-associated mutations reshape nucleic acid recognition.

### RNA asymmetrically partitions between hnRNPK and Nucleolin and selectively restrains hnRNPK condensate fusion

Having established that hnRNPK recognizes RNA in a sequence- and concentration-dependent manner at the level of the isolated protein (Fig. 2), we next interrogate whether Nucleolin engages the same 30mer MYC RNA with comparable affinity, and how the two proteins partition this shared RNA partner when co-condensed. To this end, we perform DC simulations at 310 K of hn-RNPK and Nucleolin, individually and jointly, each in the presence of 30mer MYC RNA, and quantified the resulting intermolecular contact networks and density profiles (Fig. 3). Our contact analysis—in terms of average intermolecular contact per protein residue—reveals a pronounced asymmetry in RNA binding affinity between the two proteins (Fig. 3A). In the binary mixtures, hn-RNPK engages RNA far more extensively than Nucleolin does, both in absolute terms (0.033 versus 0.009 RNA nucleotide contacts per protein residue) and relative to the RNA availability in each simulation box (∼2.4-fold versus ∼1.6-fold enrichment over a randomly mixed reference). This asymmetry is consistent with the localized interaction hotspot identified for hnRNPK (Fig. 1C), which is largely absent from Nucleolin’s more diffuse, electrostatically driven condensate interaction network. Strikingly, when hnRNPK and Nucleolin compete directly for the same RNA strands in the ternary mixture, this asymmetry does not average out but instead sharpens: the RNA enrichment of hnRNPK increases to 5-fold, while that of Nucleolin remains modest (∼1.4-fold), despite Nucleolin outnumbering hnRNPK in total residues (at the same stoichiometric molar ratio) within the simulation box. RNA therefore does not partition between the two proteins in proportion to their relative abundance, but is preferentially recruited by hnRNPK, even at Nucleolin’s expense.

**FIG. 3.**
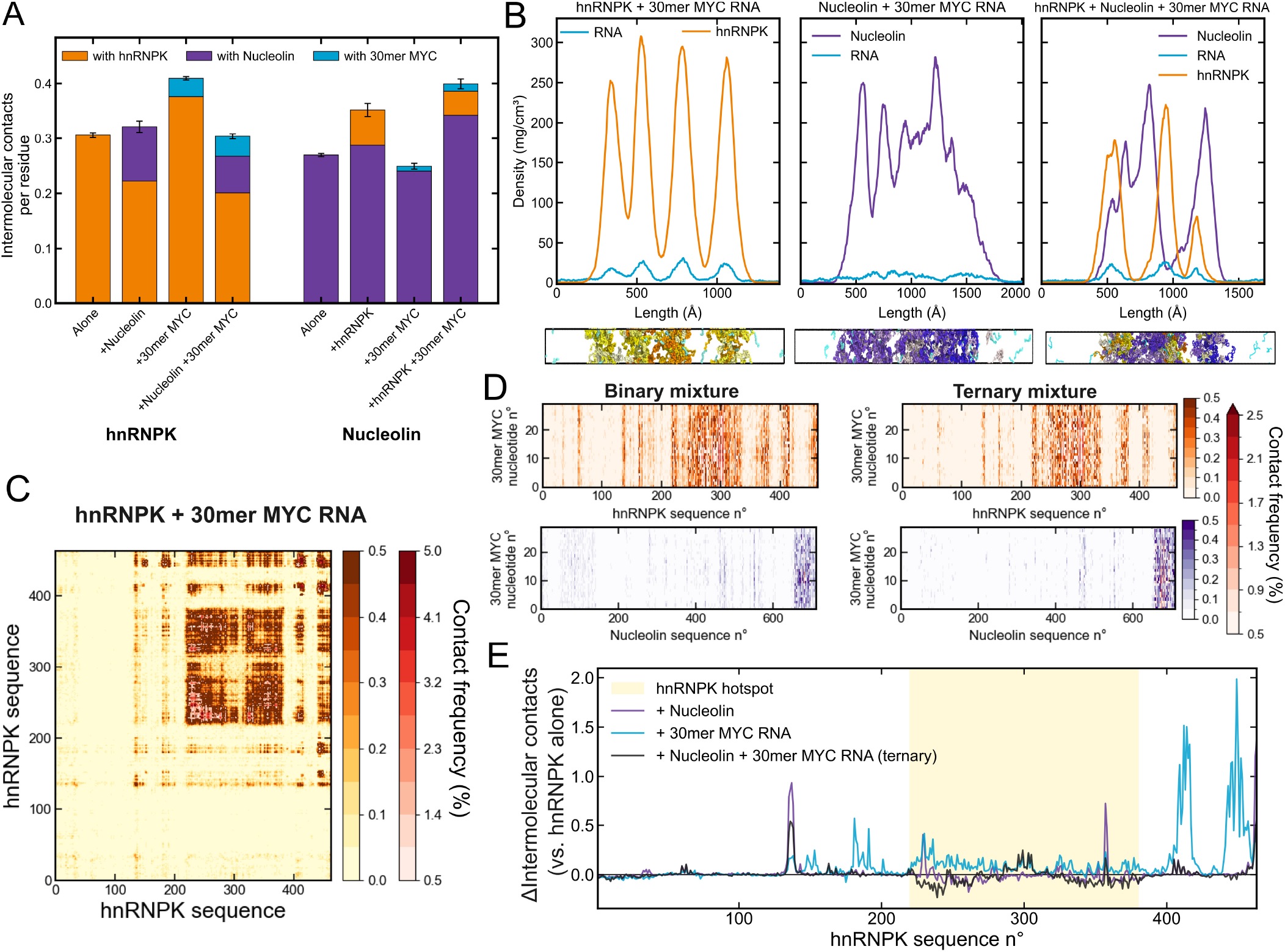
(A) Mean number of intermolecular contacts per residue displayed by hnRNPK (left) and Nucleolin (right), decomposed by contact partner (orange: hnRNPK, purple: Nucleolin, cyan: 30mer MYC RNA), for each system indicated: the homotypic protein alone, in binary mixture with the other protein, in binary mixture with RNA, and in the ternary hnRNPK–Nucleolin– RNA mixture. Values are obtained from DC simulations at 310 K by averaging over the entire trajectory; error bars denote the standard error from block averaging. (B) Density profiles along the long axis of the simulation box (top) and corresponding representative DC simulation snapshots at 310 K (bottom) for hnRNPK+30mer MYC RNA (left), Nucleolin+30mer MYC RNA (center), and the ternary hnRNPK + Nucleolin + 30mer MYC RNA mixture (right). hnRNPK replicas are colored in yellow–orange, Nucleolin chains in white–purple, and RNA chains in cyan, with individual chains distinguished by shade within each range. (C) Intermolecular contact map (in % of residue-residue contacts) of hnRNPK in the presence of 30mer MYC RNA, obtained from DC simulations of the binary mixture at 310 K. (D) Intermolecular contact maps between 30mer MYC RNA and hnRNPK (top) or Nucleolin (bottom) residues, in the binary (left) and ternary (right) mixtures at 310 K. (E) Variation in the total number of intermolecular contacts made by each hnRNPK residue, summed over all contact partners present in the system, relative to hnRNPK alone, for the two binary mixtures and the ternary mixture as indicated. The shaded region marks the hnRNPK main interaction hotspot (residues ∼220–380) identified in Fig. 1. In (A, C, D), colorbars/axes indicate the contact probability or frequency in percentage, with a two-tone scale in (C, D) distinguishing low- and high-probability contacts.

This preferential recruitment is directly visible in the spatial organization of the ternary condensate (Fig. 3B). Consistent with the partial hnRNPK–Nucleolin segregation described in Fig. 1B, the ternary density profile resolves into several protein-rich sub-regions rather than a single homogeneous slab, with the RNA density tracking the hnRNPK-rich sub-regions while remaining comparatively depleted from the Nucleolin-rich ones. This spatial coupling mirrors the behaviour of the binary systems, where the RNA density profile closely follows the hn-RNPK peaks but remains low and comparatively delocalized across the Nucleolin-rich condensate slab. Together, these results suggest that the asymmetric protein–RNA interaction grammar identified for the isolated proteins is preserved, and indeed amplified. Within the multi-component condensate, rather than being homogeneously distributed, RNA is asymmetrically partitioned toward hnRNPK-rich domains, suggesting a possible molecular mechanism by which condensate composition could locally enrich specific RNA transcripts within an otherwise mixed nucleolar-like environment^4^.

Residue-resolved contact maps reveal how this asymmetry is encoded at the sequence level. In the presence of RNA, the homotypic hnRNPK network remains essentially unperturbed (Fig. 3C), retaining the characteristic hotspot spanning residues ∼220–360 identified for the pure condensate (Fig. 1C), which indicates that RNA is accommodated within the condensate without dismantling the protein intermolecular network that sustains it. The corresponding protein–RNA maps (Fig. 3D) show that RNA engages hnRNPK densely along this same disordered region (∼220–330), with the most frequent contacts localized around residues ∼290–305, an arginine- and glycine-rich segment (∼25% Arg, ∼31% Gly) that is entirely devoid of acidic residues and therefore electrostatically optimal for binding the polyanionic RNA back-bone. Nucleolin, in contrast, contacts RNA only sparsely and almost exclusively through its C-terminal RGG domain (residues ∼645–710), the single extended basic, arginine- and glycine-rich stretch in a protein whose N-terminal half is strongly acidic (∼37% Asp/Glu and essentially devoid of arginine) and therefore electrostatically repulsive toward RNAs. This compositional contrast provides a direct sequence-level rationale for the binding asymmetry quantified in Fig. 3A, and both features become more pronounced in the ternary mixture, where the hnRNPK–RNA contact pattern consolidates within residues ∼220–340 while the already weak engagement of Nucleolin is further confined to its C-terminal RGG domain. Importantly, the residue-resolved difference of contacts between hnRNPK pure condensates and distinct mixtures (Fig. 3E) shows that RNA does not simply compete for the hnRNPK hotspot, but rather expands its interaction repertoire. Relative to hnRNPK alone, 30mer MYC RNA produces a net gain of inter-molecular contacts along most of the sequence, with the largest increases falling outside the homotypic hotspot and mapping onto the folded KH2 and KH3 domains— the canonical modules for single-stranded nucleic acid binding in hnRNPK, which contribute comparatively little to homotypic condensation. Nucleolin instead elicits far more localized changes, together with a slight depletion of contacts within the hotspot, and in the ternary mixture these opposing effects partially offset one another. hnRNPK and Nucleolin therefore recruit RNA through largely non-overlapping regions of their sequences, indicating that it is the composition-dependent redistribution of contacts, rather than a global change in interaction strength, that underlies the asymmetric organization of the ternary condensate.

Beyond its effect on the equilibrium spatial organization of the ternary condensate (Fig. 3B), RNA also exerts a marked, composition-dependent effect on condensate size dynamics (Fig. 4; cluster analysis detailed in Section SV of the SM). Tracking the number of clusters containing at least ten protein replicas over 2 *µ*s Direct Coexistence trajectories (Fig. 4A), we find that hnRNPK–RNA MYC condensates persistently fluctuate between one and four coexisting microphase-separated droplets of similar size, never settling into a single, fully fused condensate. This is directly visible in the stacked-area representation of cluster membership (Fig. 4B) and the corresponding simulation snapshots, which show hn-RNPK and RNA repeatedly redistributing across multiple, spatially separated clusters rather than coalescing into a single one. In striking contrast, Nucleolin–RNA MYC condensates rapidly converge to, and remain in, a single dominant cluster for nearly the entire trajectory, with only brief excursions to two clusters (Fig. 4A, C). RNA is therefore not merely inert toward Nucleolin at equilibrium (Fig. 3A, B), but is also dynamically agnostic to it, neither promoting nor impairing its intrinsic fusion behaviour. Notably, the ternary hnRNPK–Nucleolin– RNA mixture behaves like the Nucleolin-containing system rather than the hnRNPK-only one: it also rapidly settles into a single dominant cluster (Fig. 4A, D), suggesting that the presence of Nucleolin is sufficient to override the RNA-induced fusion impairment observed for hnRNPK in presence of 30mer MYC RNA. Together, these results reveal a second, dynamic dimension of the asymmetric protein–RNA interaction grammar identified above: RNA does not merely partition unevenly between hnRNPK and Nucleolin at equilibrium, but selectively and persistently impairs the fusion of hnRNPK-containing condensates while leaving Nucleolin-containing condensates—including the full ternary mixture—essentially unaffected.

**FIG. 4.**
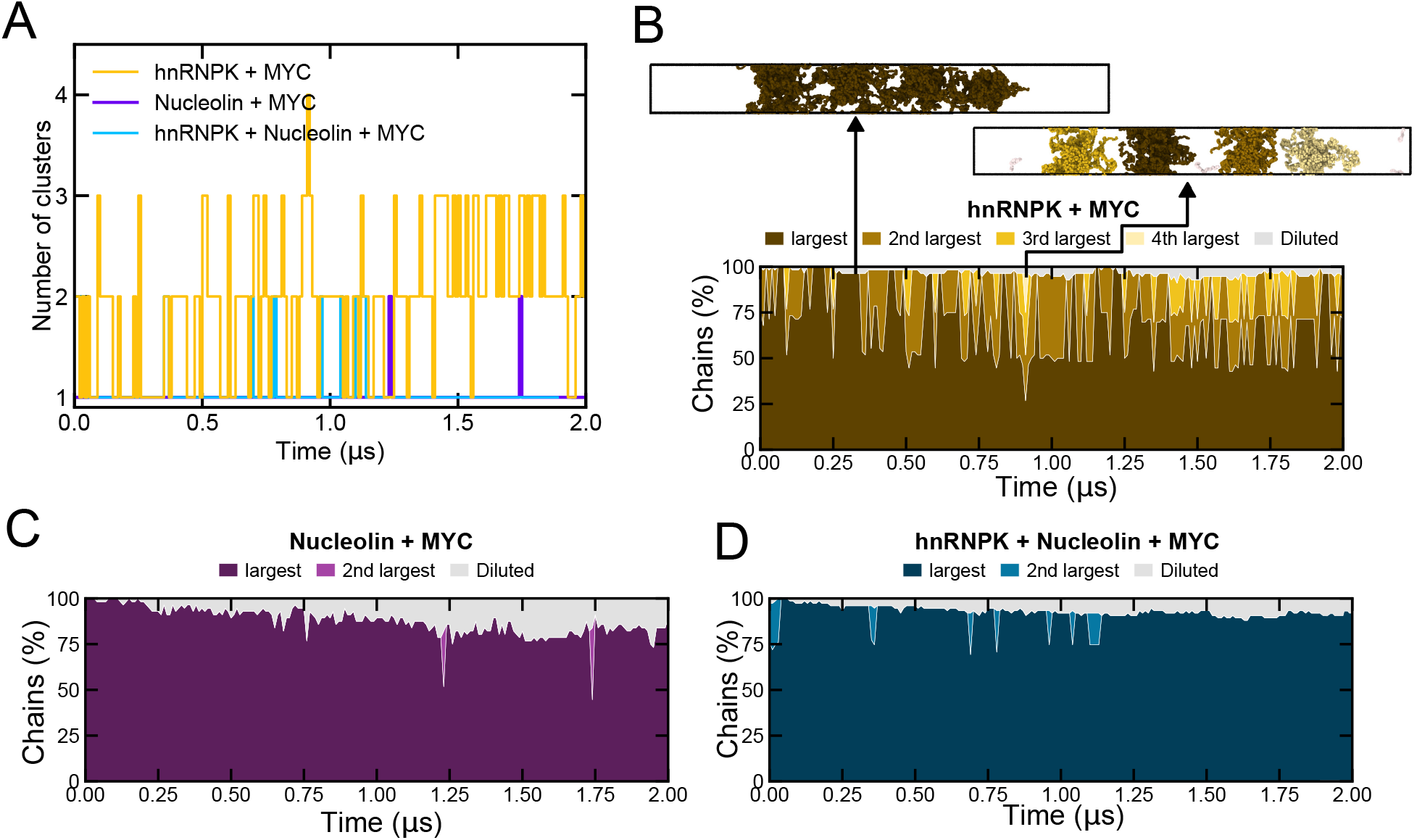
(A) Number of clusters comprising at least ten protein replicas (see Section SV of the SM) as a function of simulation time, for hnRNPK + 30mer MYC RNA (gold), Nucleolin + 30mer MYC RNA (purple), and the ternary hnRNPK + Nucleolin + 30mer MYC RNA mixture (cyan), over 2 *µ*s Direct Coexistence trajectories at 310 K. (B) Stacked-area representation of the percentage of hnRNPK chains residing in the largest, 2nd-largest, 3rd-largest, and 4th-largest clusters (dark to light gold, ranked by size at each frame) or remaining unclustered (Diluted, gray) as a function of time, for the hnRNPK + MYC RNA system. Representative simulation snapshots at the two times indicated by arrows illustrating a single, near-complete condensate (left) and a persistently fragmented state with multiple coexisting nuclei (right). Protein replicas are colored according to the size rank of the cluster they belong to at each frame (see Section SV of the SM). (C) As in (B), for the Nucleolin + MYC RNA system (largest and 2nd-largest clusters only). (D) As in (B), for the ternary hnRNPK + Nucleolin + MYC RNA mixture (largest and 2nd-largest clusters only).

## III. DISCUSSION AND CONCLUSIONS

In this work, we combine residue-resolution coarse-grained MD simulations with complementary biophysical experiments to uncover the molecular principles governing the condensation of hnRNPK and Nucleolin, two multifunctional nucleolar proteins with central roles in ribosome biogenesis and cancer biology^11–13^. By integrating quantitative phase diagrams, residue-level interaction analyses, and nucleic-acid binding experiments within a unified computational framework, we demonstrate that condensate formation cannot be understood solely from global physicochemical descriptors such as net charge or phase boundaries. Instead, our results show that condensate architecture, size, composition, and even fusion dynamics are jointly encoded in the sequence-specific organization of the intermolecular interaction network that emerges within the condensed phase—a molecular-grammar perspective that delineates the concrete mechanism by which a multiphase condensate such as the nucleolus can conserve a controlled, sub-compartmentalized architecture despite its liquid-like character^105^.

A central finding of this work is that two nucleolar proteins with a nearly indistinguishable phase diagram can nevertheless rely on fundamentally different molecular driving forces for condensation. Although hnRNPK and Nucleolin display remarkably similar critical temperatures, residue-level contact analyses reveal two distinct interaction grammars: hnRNPK condenses through a localized interaction hotspot enriched in cation–*π* and electrostatic contacts, whereas Nucleolin exploits broadly distributed long-range electrostatic interactions extending across much of its sequence. These observations reinforce the emerging view that the molecular determinants of biomolecular condensation cannot be inferred from amino acid composition alone, but instead arise from the spatial patterning of sticker-and-spacer residues along the sequence^106,107^. More broadly, they demonstrate that similar macroscopic phase behaviour may conceal fundamentally different microscopic interaction mechanisms, underscoring that sequence-resolved analyses, rather than bulk compositional metrics, are required to anticipate how a given RBP will behave once embedded in a multicomponent condensate.

The binary hnRNPK–Nucleolin mixture further shows that these interaction grammars are not fixed properties of each protein in isolation, but are context dependent. Co-condensation does not simply combine the interaction patterns of the individual proteins; instead, it produces an emergent intermolecular landscape characterized by rewiring of heterotypic contacts^84^. hnRNPK largely preserves its intrinsic interaction hotspot, whereas Nucleolin undergoes a partial redistribution of its inter-molecular contacts, replacing many homotypic electrostatic interactions with heterotypic cation–*π* and electrostatic contacts involving hnRNPK. This asymmetric response—one partner acting as a comparatively rigid interaction scaffold while the other reorganizes around it—illustrates that condensate composition actively reshapes the underlying molecular connectivity, and that multicomponent condensates cannot be interpreted as simple superpositions of their isolated constituents^108^. Such emergent, composition-dependent interaction networks likely represent a general feature of biomolecular condensates^109^, providing cells with a versatile mechanism to expand the functional and structural diversity of condensates through combinatorial assembly of different protein components, without requiring the evolution of entirely new interaction modules.

Beyond protein condensation, this work extends the Mpipi-Recharged framework to disordered single-stranded nucleic acids through a transferable, nucleotide-resolution coarse-grained RNA model compatible with residue-resolution protein simulations. Unlike previous residue-resolution treatments of protein–RNA interactions, which have generally relied on HPS-based potentials^47^, our model resolves RNA at single-nucleotide resolution with pair-specific Yukawa electrostatics, capturing both the attractive short-range interactions and the long-range electrostatic repulsion that jointly govern RNA-mediated phase separation. Consistent with this, the model successfully reproduces the experimentally observed re-entrant phase behaviour of distinct RNA–peptide condensates^26^. Although further refinements incorporating explicit counterion effects^110^ or RNA secondary structure may improve the description of highly RNA-rich systems^103^, this framework already provides a physicochemical description of experimentally relevant protein–RNA condensates, including single-stranded polyA and polyU RNAs, at a resolution that proves directly transferable to biologically realistic sequences.

We further demonstrate that the same protein– RNA interaction parameterization reproduces the concentration-dependent binding of hnRNPK to biologically relevant RUNX1 and MYC promoter-derived nucleic acid sequences. The agreement between simulation predictions and AlphaLISA experiments indicates that the model captures the underlying general physicochemical principles governing hnRNPK–nucleic acid recognition, rather than merely reproducing a specific benchmark system, and that this transferability arises naturally because the dominant interaction mechanisms—non-specific electrostatic and cation– *π* contacts—are shared by both RNA and DNA. This is of direct translational relevance: hnRNPK-mediated regulation of *RUNX*1 and *MY C* underlies its oncogenic role in acute myeloid leukemia and other hematological malignancies^12,81^, and hnRNPK has itself emerged as a druggable node, with small molecules that relocalize it away from MYC transcripts and promote its sequestration into stress granules currently being explored as a therapeutic strategy^87,89^. Therefore, a sequence-resolved computational framework for hnRNPK–nucleic acid recognition offers a route to rationalize, and potentially anticipate, how such interventions, or disease-associated mutations, may perturb this recognition process.

Beyond validating the transferability of the model, applying it to a shared RNA target reveals that RNA does not engage hnRNPK and Nucleolin equivalently, and that this asymmetry has direct consequences for both the internal organization and the dynamic behaviour of the condensates they form. RNA is preferentially and asymmetrically recruited by hnRNPK, and this asymmetry sharpens, rather than averaging out, when the two proteins compete for the same RNA pool within a ternary condensate, spatially partitioning RNA toward hnRNPK-rich sub-regions—reminiscent of how competing protein–RNA interaction networks have been proposed to organize immiscible sub-phases within multi-phase condensates more generally^85^. Strikingly, this compositional asymmetry extends to condensate dynamics: cluster analysis of the simulated trajectories shows that RNA selectively and persistently impairs the fusion rate and condensate size of hnRNPK-containing condensates, which remain fragmented into multiple, unfused clusters over microsecond timescales. In contrast, Nucleolin-containing condensates—including the full ternary mixture—coalesce into a single dominant cluster largely unaffected by 30mer MYC RNAs. This dual effect on both condensate size and fusion kinetics identifies RNA as a composition-selective regulator that can simultaneously enrich specific sub-compartments with RNAs and restrain their coalescence. This offers a plausible mechanism by which a multiphase condensate such as the nucleolus—long known to comprise coexisting, immiscible liquid sub-phases^105^—could conserve multiple, compositionally distinct sub-regions of controlled size rather than collapsing into a single, large, well-mixed droplet.

Together, these results support a model in which the sequence-specific interplay between hnRNPK, Nucleolin, and RNA does not merely dictate whether these components phase separate, but actively shapes the internal organization, compositional identity, and fusion dynamics of the condensates they form—properties that are central to how a condensate such as the nucleolus establishes and conserves its characteristic size and multiphase architecture over time. Because these behaviours emerge from residue-level sequence features rather than bulk physicochemical descriptors, this modelling framework provides a quantitative platform for predicting how sequence variation, post-translational modifications, or disease-associated mutations affecting hnRNPK, Nucleolin, or their RNA/RBP partners might perturb this organization. Such perturbations are far from hypothetical, since hnRNPK dysregulation is directly implicated in acute myeloid leukemia and other ribosomopathy-like phenotypes^11–13^, and altered nucleolar number, size, and morphology remain recurrent hallmarks of cancer, ribosomopathies, and ageing more broadly. We propose a computational residue-level approach, which combined with stringent biochemical validation, may help guide systematic investigation of the molecular mechanisms underlying nucleolar dysfunction, and more generally of how sequence-encoded protein–RNA interaction networks tune the assembly, organization, and fusion dynamics of multiphase condensates. Beyond nucleolar organization, hnRNPK condensates have recently been shown to facilitate enhancer–promoter looping and the recruitment of RNA polymerase II^111^, raising the possibility that the sequence-encoded interaction network characterized here also feeds into transcriptional regulation; addressing this directly will require simulations of hnRNPK/Nucleolin condensates together with RNA polymerase II components, which we leave for future work.

## IV. DATA AVAILABILITY

All data supporting the findings of this study are available within the article and its Supplementary Data. The complete set of coarse-grained RNA model parameters (Wang–Frenkel and Yukawa interaction parameters for all RNA–RNA and RNA–protein pairs) is provided in Section SIV of the SM. Simulation input files, force-field files, representative trajectories and the analysis scripts used to generate all figures are available from the corresponding author upon reasonable request.

## V. ACKNOWLEDGMENTS

The authors acknowledge the computational resources provided by the Red Española de Supercomputación (RES) at the Barcelona Supercomputing Center (BSC), through projects FI-2025-3-0003, FI-2025-3-0065 and EHPC-REG-2025R02-173 on the MareNostrum5 supercomputer, and the computational resources at the CIEMAT Xula supercomputer through project FI-2026-1-0027.

