## Supplementary Material for "Size Control of hnRNPK-based Nucleolar Condensates by RNA-Regulated Fusion Dynamics"

Maria Velasco-Estevez

*H12O-CNIO Hematological Malignancies Clinical Research Unit. Spanish Cancer Research Center (CNIO), Madrid*

Alberto Ocana

*Experimental Therapeutics in Cancer Unit, Instituto de Investigación Sanitaria San Carlos (IdISSC), and CIBERONC, Madrid, Spain and PhAsIca Biosciences S.L, Calle Velázquez, 27, 28001 Madrid, Spain*

Rosana Collepardo

*Yusuf Hamied Department of Chemistry, University of Cambridge, Lensfield Road, Cambridge CB2 1EW, UK Department of Genetics, University of Cambridge,*

*Cambridge CB2 3EH, United Kingdom.*
*Maxwell Centre, Cavendish Laboratory, Department of Physics,*
*University of Cambridge, J J Thomson Avenue,*
*Cambridge CB3 0HE, United Kingdom. and*
*PhAsIca Biosciences S.L, Calle Velázquez, 27, 28001 Madrid, Spain*

Jorge R. Espinosa*
*Department of Physical Chemistry, Universidad Complutense de Madrid,*
*Av. Complutense s/n, Madrid 28040, Spain*
*Instituto Pluridisciplinar, Universidad Complutense de Madrid,*
*P.º de Juan XXIII, 1, Moncloa - Aravaca, 28040 Madrid, Spain*
*Yusuf Hamied Department of Chemistry, University of Cambridge,*
*Lensfield Road, Cambridge CB2 1EW, UK and*
*PhAsIca Biosciences S.L, Calle Velázquez, 27, 28001 Madrid, Spain*

(Dated: September 11, 2026)

*

### SI. THE MPIPI-RECHARGED MODEL

The Mpipi-Recharged model is a residue-level coarse-grained force field for both protein and protein/RNA condensates [1]. Each amino acid or nucleotide is represented by a single bead connected to adjacent residues by harmonic bonds. Globular domains of the proteins are treated as rigid bodies whose beads are fixed at the $C_\alpha$ position from the corresponding Protein Data Bank (PDB). The potential energy is computed as the sum of pairwise bonded ($E_{\text{bonded}}$) and non-bonded ($E_{\text{non-bonded}}$) interactions as:

$$E = E_{\text{bonded}} + E_{\text{non-bonded}}. \tag{S1}$$

The intrinsically disordered regions (IDRs) are modeled as fully flexible polymers and these are connected to the globular domains. The bonded potential is written as:

$$E_{\text{bonded}}(r_{ij}) = \sum_{ij} k(r_{ij} - r_0)^2, \tag{S2}$$

where $r_{ij}$ is the distance between the connected beads, $r_0 = 3.81\,\text{Å}$ and $r_0 = 5.00\,\text{Å}$ are the equilibrium bond lengths for protein and RNA, respectively. The spring constant $k = 9.6\,\text{kcal}\cdot\text{mol}^{-1}\cdot\text{Å}^{-2}$. The sum runs over all bonded residues.

Non-bonded interactions consist of the sum of the hydrophobic interaction and the electrostatic interaction. The hydrophobic interaction is given by the Wang–Frenkel (WF) potential [2] that accounts for short-ranged excluded-volume repulsion and long-ranged attraction. This potential is defined as:

$$E_{\text{WF}}(r_{i,j}) = \sum_{ij} \epsilon_{ij}\alpha_{ij} \left[\left(\frac{\sigma_{ij}}{r_{ij}}\right)^{2\mu_{ij}} - 1\right] \left[\left(\frac{R_{ij}}{r_{ij}}\right)^{2\mu_{ij}} - 1\right]^{2\nu_{ij}}, \tag{S3}$$

where

$$\alpha_{ij} = 2\nu_{ij}\left(\frac{R_{ij}}{\sigma_{ij}}\right)^{2\mu_{ij}} \left\{\frac{2\nu_{ij}+1}{2\nu_{ij}\left[\left(\frac{R_{ij}}{\sigma_{ij}}\right)^{2\mu_{ij}} - 1\right]}\right\}^{2\nu_{ij}+1}. \tag{S4}$$

Here $\sigma_{ij}$ is the pair-of-beads diameter, defined from the individual diameters ($\sigma_i$ and $\sigma_j$) assuming the Lorentz-Berthelot mixing rules (i.e., $\sigma_{ij} = (\sigma_i + \sigma_j)/2$). $R_{ij} = 3\sigma_{ij}$ is the cut-off distance for the $ij$-th interaction. The interaction parameter $\epsilon_{ij}$ is defined for each specific amino acid pair based on our atomistic Potential of Mean Force calculations and bioinformatics data [1]. The exponent $\nu_{ij}$ is set to 1 for all pairs and $\mu_{ij}$ depends on the specific pair, ranging from 2 to 12 (see Ref. [1]). Notice that higher values of $\mu_{ij}$ lead to a steeper increase in the repulsive part of the potential. The interaction involving globular domains are reduced to account for the 'buried' interactions. In particular, the interaction between flexible regions and globular domains are screened by a factor of $\sqrt{0.7}$ and the WF interaction between residues in globular regions is scaled down a factor of 0.7.

Electrostatic interactions are described by the Yukawa potential [3] instead of the Debye–Hückel potential [4] originally employed in the Mpipi model. Avoiding the use of explicit charge values, this potential allows to modulate independently the strength of electrostatic interactions in a pair-specific basis. This potential is defined as:

$$E_{\text{electrostatic}} = \sum_{ij} \frac{A_{ij}}{r_{ij}} \exp(-\kappa r_{ij}), \qquad r_{ij} < r_c, \tag{S5}$$

where $A_{ij}$ is the interaction parameter, $\kappa$ is the screening due to ions, and $r_{ij}$ is the distance between interacting residues. The cut-off for the electrostatic interaction, $r_c$, is set to 3.5 nm. The screening parameter $\kappa$ is expressed in a explicit way as $\kappa = \sqrt{8\pi B c_s}$, where $c_s$ is the salt concentration—normally set to 150 mM of NaCl—and $B = e_0^2/4\pi k_B T \varepsilon_0 \varepsilon_r$ is the Bjerrum length. The relative dielectric constant $\varepsilon_r$ varies with temperature according to the empiric formula [5]:

$$\varepsilon_r(T) = \frac{5321}{T} + 233.760 - 0.9297T + 1.417 \cdot 10^{-3} T^2 - 8.292 \cdot 10^{-7} T^3, \tag{S6}$$

for $T$ in Kelvin. The pair-specific optimized values of $A_{ij}$ of the Yukawa potential can be found in Ref. [1]. In the Mpipi-Recharged model, such parameters indicate that the interaction between oppositely charged pairs is significantly stronger than those between identically charged pairs.

### SII. SEQUENCES AND PDBS OF THE STUDIED PROTEINS

The globular domains for hnRNPK and Nucleolin was extracted from the AF prediciton considering Globular when the prediction is above 0.9 of pLDDT.

**hnRNPK:**

```
METEQPEETFPNTETNGEFGKRPAEDMEEEQAFKRSRNTDEMVELRILLQSKNAGAVIGKGGKNIKALRTDYNASVSV
PDSSGPERILSISADIETIGEILKKIIPTLEEGLQLPSPTATSQLPLESDAVECLNYQHYKGSDFDCELRLLIHQSLA
GGIIGVKGAKIKELRENTQTTIKLFQECCPHSTDRVVLIGGKPDRVVECIKIILDLISESPIKGRAQPYDPNFYDETY
DYGGFTMMFDDRRGRPVGFPMRGRGGFDRMPPGRGGRPMPPSRRDYDDMSPRRGPPPPPPGRGGRGGSRARNLPLPPP
PPPRGGDLMAYDRRGRPGDRYDGMVGFSADETWDSAIDTWSPSEWQMAYEPQGGSGYDYSYAGGRGSYGDLGGPIITT
QVTIPKDLAGSIIGKGGQRIKQIRHESGASIKIDEPLEGSEDRIITITGTQDQIQNAQYLLQNSVKQYSGKFF
```

**Nucleolin:**

```
MVKLAKAGKNQGDPKKMAPPPKEVEEDSEDEEMSEDEEDDSSGEEVVIPQKKGKKAAATSAKKVVVSPTKKVAVATPA
KKAAVTPGKKAAATPAKKTVTPAKAVTTPGKKGATPGKALVATPGKKGAAIPAKGAKNGKNAKKEDSDEEEDDDSEED
EEDDEDEDEDEDEIEPAAMKAAAAAPASEDEDDEDDEDDEDDDDDEEDDSEEEAMETTPAKGKKAAKVVPVKAKNVAE
DEDEEEDDEDEDDDDDEDDEDDDDEDDEEEEEEEEEEEPVKEAPGKRKKEMAKQKAAPEAKKQKVEGTEPTTAFNLFVG
NLNFNKSAPELKTGISDVFAKNDLAVVDVRIGMTRKFGYVDFESAEDLEKALELTGLKVFGNEIKLEKPKGKDSKKER
DARTLLAKNLPYKVTQDELKEVFEDAAEIRLVSKDGKSKGIAYIEFKTEADAEKTFEEKQGTEIDGRSISLYYTGEKG
QNQDYRGGKNSTWSGESKTLVLSNLSYSATEETLQEVFEKATFIKVPQNQNGKSKGYAFIEFASFEDAKEALNSCNKR
EIEGRAIRLELQGPRGSPNARSQPSKTLFVKGLSEDTTEETLKESFDGSVRARIVTDRETGSSKGFGFVDFNSEEDAK
AAKEAMEDGEIDGNKVTLDWAKPKGEGGFGGRGGGRGGFGGRGGGRGGRGGFGGRGRGGFGGRGGFRGGRGGGGDHKP
QGKKTKFE
```

The RNA sequences simulated along with hnRNPK and Nucleolin are presented below (uracil, U, replacing thymine, T, relative to the DNA constructs used in the AlphaLISA experiments).

**30mer MYC:**

```
CGACCCCUCG GUGGUCUUCC CCUACCCUC
```

**28mer RUNX1:**

```
CGCCCCCCCC CACCCCCCGC
```

**38mer RUNX1:**

```
CGCCCCCCCC CACCCCCCGC AGUAAUAAAG GCCCCUGA
```

### SIII. CONTACT MAPS

Intermolecular contact maps were computed from the Direct Coexistence trajectories by counting, for every residue pair $(i, j)$ belonging to different chains, the number of frames in which the distance between beads $i$ and $j$ was below a cutoff of $1.2\,\sigma_{ij}$, averaged over all sampled chain pairs and simulation frames. The resulting contact probability (in %) is reported as a function of sequence position for the homotypic and heterotypic maps discussed in the main text.

To identify the specific residue-residue interactions underlying the heterotypic hnRNPK–

Nucleolin contact map (Figure 1E of the main text), we further decomposed the ten most frequent residue-pair contacts by amino-acid identity, normalized to the most frequent pair. Fig. S1 shows this decomposition, with contacts colored by interaction type: charged–charged in blue, cation–$\pi$ in pink, $\pi$–$\pi$ in green, and all other contact types in gray.

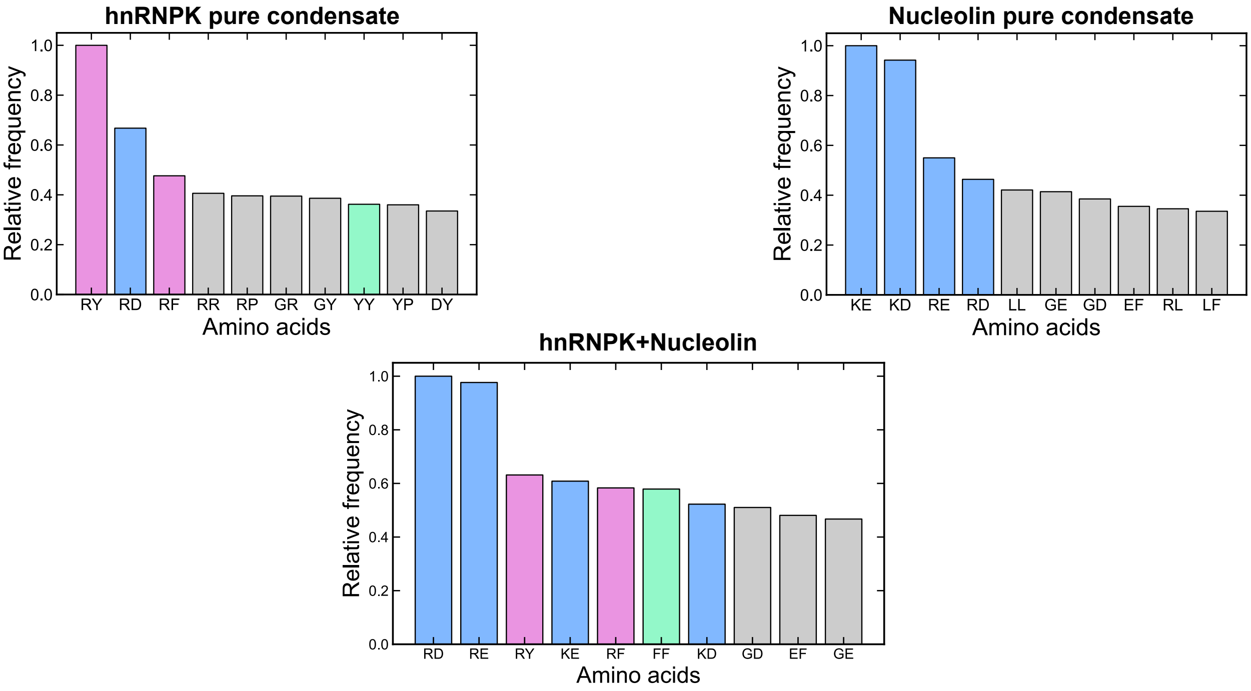


FIG. S1. Ten most frequent residue-residue contact pairs between hnRNPK in pure condensates, Nucleolin in pure condensates, and hnRNPK-Nucleolin in the equimolar binary mixture at 310 K, as indicated, obtained from the heterotypic contact map (Figure 1E of the main text), ranked and normalized by relative frequency. Charged–charged interactions are coloured in blue, cation–$\pi$ in pink, $\pi$–$\pi$ in green, and all other contact types in gray.

For the equimolar hnRNPK–Nucleolin mixture, the homotypic hnRNPK–hnRNPK and Nucleolin–Nucleolin contact maps are shown in Figs. S2 and S3, respectively, for comparison with those of the corresponding single-component condensates (Figures 1C and 1D of the main text).

### SIV. MPIPI-RECHARGED PARAMETERS

The electrostatic interactions in the Mpipi-Recharged model are described by the Yukawa potential introduced in Eq. S5, with a pair-specific interaction parameter $A_{ij}$. The analogous Coulomb potential is given by

$$E_{Coulomb} = \frac{k}{\epsilon}\frac{q_i q_j}{r}, \tag{S7}$$

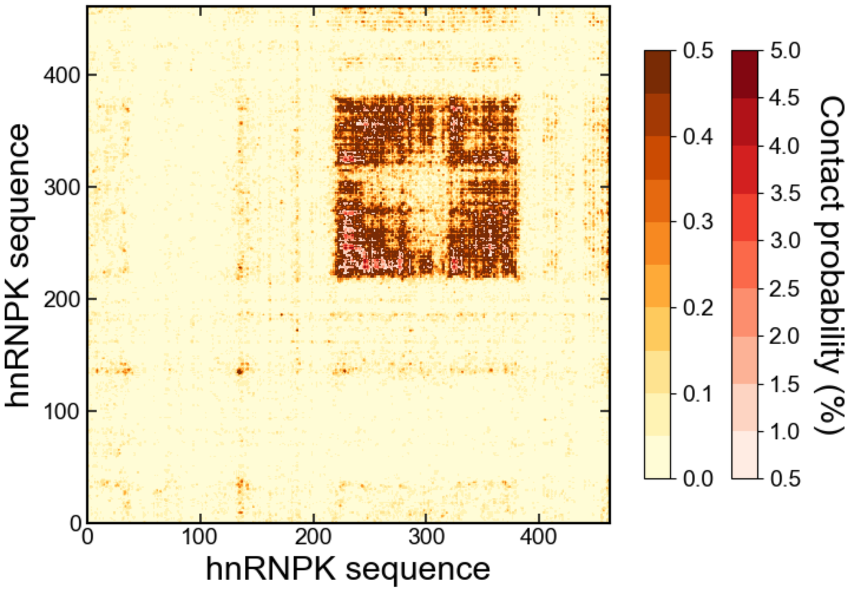


FIG. S2. Homotypic hnRNPK–hnRNPK intermolecular contact map in the equimolar hnRNPK–Nucleolin condensate at 310 K. Contact probabilities from 0 to 0.5% are shown in the orange scale and from 0.5 to 5% in the red scale.

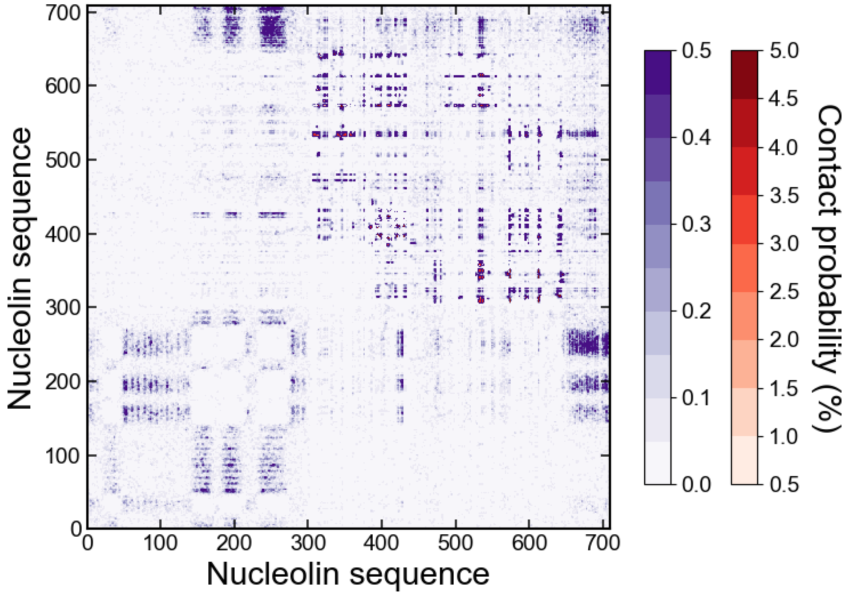


FIG. S3. Homotypic Nucleolin–Nucleolin intermolecular contact map in the equimolar hnRNPK–Nucleolin condensate at 310 K. Contact probabilities from 0 to 0.5% are shown in the purple scale and from 0.5 to 5% in the red scale.

where $k = 1/4\pi\varepsilon_0$ is the Coulombic constant, $\epsilon$ is the dielectric constant of the medium, and $q_i$, $q_j$ are the charges of the interacting particles. Given this potential, the interaction parameter for the Yukawa potential is calculated as $A_{ij} = k\, q_i q_j/\epsilon$. For the values typically used in previous models, *i.e.* $k = 331.61$ kcal mol$^{-1}e^{-2}$ Å, $\epsilon = 80$, and $q_{i,j} = \pm 1e$ (or $q_H = +0.5e$ for histidine), the magnitude of the interaction is $|A_{ij}| = 4.145$ (or $|A_{iH}| = 2.073$ when involving histidine). The sign of $A_{ij}$ depends on whether the interaction is attractive or repulsive; the parameter $A_{ij}$ can therefore be directly converted into the product $q_i q_j$ to recover the effective charge product being modeled by a Coulomb-like potential.

The complete set of protein–protein short-range (Wang–Frenkel) and electrostatic (Yukawa) parameters of the Mpipi-Recharged model is given in Ref. [1] and is not reproduced here; Section SIV A below provides the corresponding RNA–RNA and RNA–protein parameters introduced in this work.

#### A. RNA and RNA–protein parameters

The RNA nucleotide-nucleotide and nucleotide-amino acid short-range (Wang–Frenkel) parameters are given in Table S1 (RNA–RNA) and Table S2 (RNA–protein), following the same Wang–Frenkel functional form used for the protein–protein interactions of the original Mpipi-Recharged model [1].

| Nucleotide i$^{\text{th}}$ | Nucleotide j$^{\text{th}}$ | $\epsilon$ (kcal mol$^{-1}$) | $\sigma$ (Å) | $\nu$ | $\mu$ |
|---|---|---|---|---|---|
| **A** | **A** | 0.0850 | 8.4400 | 1 | 4 |
| **A** | **C** | 0.1082 | 8.3300 | 1 | 4 |
| **A** | **G** | 0.0398 | 8.4750 | 1 | 7 |
| **A** | **U** | 0.1408 | 8.3050 | 1 | 3 |
| **C** | **C** | 0.1250 | 8.2200 | 1 | 4 |
| **C** | **G** | 0.1620 | 8.3650 | 1 | 3 |
| **C** | **U** | 0.0702 | 8.1950 | 1 | 4 |
| **G** | **G** | 0.0957 | 8.5100 | 1 | 3 |
| **G** | **U** | 0.1202 | 8.3400 | 1 | 3 |
| **U** | **U** | 0.1102 | 8.1700 | 1 | 3 |

TABLE S1: RNA–RNA short-range (Wang–Frenkel) interaction parameters of the Mpipi-Recharged RNA model.

| Amino acid | Nucleotide | $\epsilon$ (kcal mol$^{-1}$) | $\sigma$ (Å) | $\nu$ | $\mu$ |
|---|---|---|---|---|---|
| **M** | **A** | 0.1140 | 7.4540 | 1 | 4 |
| **M** | **C** | 0.0895 | 7.3440 | 1 | 4 |
| **M** | **G** | 0.1251 | 7.4890 | 1 | 4 |
| **M** | **U** | 0.1723 | 7.3190 | 1 | 3 |
| **G** | **A** | 0.3687 | 6.5676 | 1 | 3 |
| **G** | **C** | 0.2330 | 6.4576 | 1 | 3 |
| **G** | **G** | 0.3549 | 6.6026 | 1 | 3 |
| **G** | **U** | 0.2007 | 6.4326 | 1 | 3 |
| **K** | **A** | 0.0924 | 7.5557 | 1 | 4 |
| **K** | **C** | 0.1511 | 7.4457 | 1 | 4 |
| **K** | **G** | 0.1257 | 7.5907 | 1 | 4 |
| **K** | **U** | 0.0972 | 7.4207 | 1 | 3 |
| **T** | **A** | 0.1079 | 7.1645 | 1 | 4 |
| **T** | **C** | 0.0442 | 7.0545 | 1 | 4 |
| **T** | **G** | 0.1098 | 7.1995 | 1 | 4 |
| **T** | **U** | 0.1679 | 7.0295 | 1 | 3 |
| **R** | **A** | 0.4908 | 7.6395 | 1 | 3 |
| **R** | **C** | 0.3197 | 7.5295 | 1 | 3 |
| **R** | **G** | 0.3596 | 7.6745 | 1 | 3 |
| **R** | **U** | 0.3949 | 7.5045 | 1 | 3 |
| **A** | **A** | 0.2468 | 6.8550 | 1 | 3 |
| **A** | **C** | 0.1984 | 6.7450 | 1 | 3 |
| **A** | **G** | 0.1984 | 6.8900 | 1 | 3 |
| **A** | **U** | 0.1772 | 6.7200 | 1 | 3 |
| **D** | **A** | 0.1425 | 7.1318 | 1 | 3 |
| **D** | **C** | 0.2807 | 7.0218 | 1 | 3 |
| **D** | **G** | 0.2368 | 7.1668 | 1 | 3 |
| **D** | **U** | 0.1920 | 6.9968 | 1 | 3 |

| | | | | | |
|---|---|---|---|---|---|
| **E** | **A** | 0.2610 | 7.3088 | 1 | 3 |
| **E** | **C** | 0.1065 | 7.1988 | 1 | 3 |
| **E** | **G** | 0.3416 | 7.3438 | 1 | 3 |
| **E** | **U** | 0.1953 | 7.1738 | 1 | 3 |
| **Y** | **A** | 0.5800 | 7.5868 | 1 | 3 |
| **Y** | **C** | 0.5879 | 7.4768 | 1 | 3 |
| **Y** | **G** | 0.8084 | 7.6218 | 1 | 3 |
| **Y** | **U** | 0.6621 | 7.4518 | 1 | 3 |
| **V** | **A** | 0.1313 | 7.3530 | 1 | 4 |
| **V** | **C** | 0.0979 | 7.2430 | 1 | 4 |
| **V** | **G** | 0.1044 | 7.3880 | 1 | 4 |
| **V** | **U** | 0.1553 | 7.2180 | 1 | 3 |
| **L** | **A** | 0.1173 | 7.4870 | 1 | 4 |
| **L** | **C** | 0.0910 | 7.3770 | 1 | 4 |
| **L** | **G** | 0.0984 | 7.5220 | 1 | 4 |
| **L** | **U** | 0.1580 | 7.3520 | 1 | 3 |
| **Q** | **A** | 0.1167 | 7.3589 | 1 | 4 |
| **Q** | **C** | 0.3025 | 7.2489 | 1 | 4 |
| **Q** | **G** | 0.3019 | 7.3939 | 1 | 4 |
| **Q** | **U** | 0.2527 | 7.2239 | 1 | 3 |
| **W** | **A** | 0.5208 | 7.7533 | 1 | 3 |
| **W** | **C** | 0.3604 | 7.6433 | 1 | 3 |
| **W** | **G** | 0.4027 | 7.7883 | 1 | 3 |
| **W** | **U** | 0.4276 | 7.6183 | 1 | 3 |
| **F** | **A** | 0.3627 | 7.5348 | 1 | 3 |
| **F** | **C** | 0.3303 | 7.4248 | 1 | 3 |
| **F** | **G** | 0.3494 | 7.5698 | 1 | 3 |
| **F** | **U** | 0.3483 | 7.3998 | 1 | 3 |
| **S** | **A** | 0.0936 | 6.9263 | 1 | 4 |

| | | | | | |
|---|---|---|---|---|---|
| **S** | **C** | 0.0786 | 6.8163 | 1 | 4 |
| **S** | **G** | 0.1634 | 6.9613 | 1 | 4 |
| **S** | **U** | 0.1833 | 6.7913 | 1 | 3 |
| **H** | **A** | 0.1112 | 7.3889 | 1 | 4 |
| **H** | **C** | 0.4432 | 7.2789 | 1 | 4 |
| **H** | **G** | 0.4566 | 7.4239 | 1 | 4 |
| **H** | **U** | 0.1661 | 7.2539 | 1 | 3 |
| **N** | **A** | 0.1886 | 7.1817 | 1 | 3 |
| **N** | **C** | 0.2812 | 7.0717 | 1 | 3 |
| **N** | **G** | 0.2911 | 7.2167 | 1 | 3 |
| **N** | **U** | 0.2494 | 7.0467 | 1 | 3 |
| **P** | **A** | 0.2509 | 7.1231 | 1 | 3 |
| **P** | **C** | 0.2122 | 7.0131 | 1 | 3 |
| **P** | **G** | 0.2776 | 7.1581 | 1 | 3 |
| **P** | **U** | 0.1918 | 6.9881 | 1 | 3 |
| **C** | **A** | 0.1425 | 7.0822 | 1 | 3 |
| **C** | **C** | 0.0939 | 6.9722 | 1 | 3 |
| **C** | **G** | 0.1714 | 7.1172 | 1 | 3 |
| **C** | **U** | 0.1872 | 6.9472 | 1 | 3 |
| **I** | **A** | 0.1148 | 7.6808 | 1 | 4 |
| **I** | **C** | 0.0887 | 7.5708 | 1 | 4 |
| **I** | **G** | 0.0957 | 7.7158 | 1 | 4 |
| **I** | **U** | 0.1527 | 7.5458 | 1 | 3 |

TABLE S2: RNA–protein short-range (Wang–Frenkel) interaction parameters of the Mpipi-Recharged RNA model.

For the electrostatic (Yukawa) interactions, all ten RNA–RNA nucleotide pairs share the same parameter, $A_{ij} = +4.1516$ kcal mol$^{-1}$ Å (equivalent to $q_i q_j = +1$, since every nucleotide carries the same unit negative charge on its phosphate backbone), reflecting the uniform electrostatic repulsion between RNA strands. The pair-specific values for the electrostatic interaction between the charged/polar amino acids and each of the four ribonucleotides are given in Table S3, following the general $A_{ij} \leftrightarrow q_i q_j$ correspondence described above.

| | **A** | **C** | **G** | **U** |
|---|---|---|---|---|
| **K** | -4.34 | -4.24 | -4.24 | -4.1516 |
| **R** | -4.92 | -4.72 | -4.81 | -4.1516 |
| **D** | +4.00 | +4.00 | +4.00 | +4.1516 |
| **E** | +4.00 | +4.00 | +4.00 | +4.1516 |
| **H** | -2.17 | -2.17 | -2.17 | -2.0758 |

TABLE S3. Parameter $A_{ij}$ (in kcal mol$^{-1}$ Å) for the Yukawa potential between each charged/polar amino acid (rows) and each ribonucleotide (columns).

### SV. CONDENSATE CLUSTER ANALYSIS

To quantify condensate formation and its evolution over the course of the Direct Coexistence simulations, we performed a cluster analysis of the corresponding trajectories. Each simulated chain was assigned to a species (hnRNPK, Nucleolin, or RNA) based on its bead composition and length, and two chains were considered to be in contact, and therefore part of the same cluster, if they shared at least two simultaneous inter-chain bead–bead contacts within 25 Å of one another. This distance is not an arbitrary choice: it corresponds to the short-range interaction cutoff of the Wang–Frenkel potential used in the Mpipi-Recharged force field itself, so that the clustering criterion reflects the same interaction range that drives condensation in the simulation. A criterion based on the whole-chain center-of-mass distance was also considered, but discarded, as it requires an arbitrary cutoff that becomes unreliable when chains of substantially different size coexist in the same system, as is the case for hnRNPK and Nucleolin. Requiring at least two simultaneous contacts, rather than a single one, removed spurious short-lived associations arising from transient individual bead contacts without altering the identity or behaviour of the dominant condensate; increasing the threshold to three contacts gave essentially indistinguishable results, confirming that the analysis is converged with respect to this choice.

Clusters were defined as the connected components of the resulting chain–chain contact network, obtained via a standard union–find algorithm. A connected component was counted as a genuine condensate for the purposes of tracking the number of clusters and the cluster membership of individual chains over time only once it comprised at least ten chains; smaller connected components were treated as sub-threshold oligomers rather than condensates. This size threshold was not applied when reporting the size or molecular composition of the single largest cluster, nor the overall distribution of cluster sizes, which were computed for all connected components regardless of size.

For each trajectory we report, as a function of simulation time: the number of qualifying clusters; the size of the single largest cluster; and the molecular composition of the largest cluster (i.e., the number of chains of each species it contains), which indicates whether the dominant condensate is a genuine multicomponent assembly rather than one species aggregating separately nearby. We additionally report the time-averaged distribution of cluster sizes. To visualize how individual chains partition among condensates over time, each chain was assigned, at every frame, a color corresponding to the size rank of the cluster it belonged to at that instant (e.g., the largest cluster, the second largest, and so on), rather than to the identity of a specific, persistent condensate. This rank-based convention avoids the ambiguity that arises when a condensate splits into two or more daughter clusters, which cannot be unambiguously assigned to a single parent identity.

### SVI. ALPHALISA EXPERIMENTAL ASSAY

To experimentally validate the binding of hnRNPK to the MYC and RUNX1 sequences, we used AlphaLISA technology (PerkinElmer). Full-length His-tagged hnRNPK (FL-hnRNPK) protein was purified following recombinant protein expression in *E. coli* BL21 cells, in accordance with the protocol pipeline established by the CNIO Crystallography Unit Service [6]. Biotinylated single-stranded DNA (ssDNA) oligonucleotides corresponding to both the MYC and RUNX1 sequences were custom-designed and purchased from Sigma-Aldrich.

For the AlphaLISA binding assay, streptavidin donor beads (#6760002, PerkinElmer) and anti-6xHis acceptor beads (AL178M, PerkinElmer) were used to tag the ssDNA and hnRNPK, respectively, at a 1:1 bead ratio. The concentration of hnRNPK was kept constant at 7.5 $\mu\mathrm{g\,ml^{-1}}$ while the concentration of ssDNA was titrated. Fluorescence was measured using an EnVision 2104 Multilabel Reader (PerkinElmer) with excitation at 680 nm and emission at 560 nm, following the manufacturer's instructions.

---
